# Dimerization of iLID Optogenetic Proteins Observed Using 3D Single-Molecule Tracking in Live Bacterial Cells

**DOI:** 10.1101/2022.07.10.499479

**Authors:** Alecia M. Achimovich, Ting Yan, Andreas Gahlmann

**Author notes:** Address correspondence to Andreas Gahlmann.

## Abstract

3D single molecule tracking microscopy has enabled measurements of protein diffusion in living cells, offering information about protein dynamics and cellular environments. For example, different diffusive states can be resolved and assigned to protein complexes of different size and composition. However, substantial statistical power and biological validation, often through genetic deletion of binding partners, are required to support diffusive state assignments. When investigating cellular processes, transient perturbation to protein spatial distributions is preferable to permanent genetic deletion of an essential protein. In this context, optogenetic dimerization systems can be used to manipulate protein spatial distributions which could offer a means to deplete specific diffusive states observed in single molecule tracking experiments. Here, we evaluate the performance of the iLID optogenetic system in living *E. coli* cells using diffraction-limited microscopy and 3D single-molecule tracking. We observed a robust optogenetic response in protein spatial distributions after 488 nm laser activation. Surprisingly, 3D single-molecule tracking results indicate activation of the optogenetic response when illuminating with high intensity light with wavelengths at which there is minimal photon absorbance by the LOV2 domain. The pre-activation can be minimized through the use of iLID system mutants, and titration of protein expression levels.

**SIGNIFICANCE STATEMENT:** We describe the combination of 3D single-molecule tracking microscopy and optogenetic manipulation of protein spatial distributions in living cells. Such a combination is impactful, because optogenetic systems enable sample perturbation in real time using light, which provides more flexibility than gene deletion and gene editing approaches that result in permanent changes to the specimen. We specifically investigate the performance of the iLID optogenetic system in a knocksideways experiment, in which cytosolic prey proteins (SspB) are sequestered to the membrane by interacting with membrane-anchored bait proteins (iLID). We quantified the magnitude of the optogenetic effect using both diffraction-limited imaging and 3D single-molecule tracking microscopy. We found, surprisingly, that the iLID optogenetic response is activated substantially by high intensity light at wavelengths for which there is negligible photon absorption by the iLID protein. Quantification of this alternative activation mechanism is a necessary component before optogenetic tools, such as iLIDs, are employed in single-molecule knocksideways experiment that are designed to provide new biological insights.

## INTRODUCTION

Many fluorescence microscopy-based techniques have been developed over the past decades to detect and quantify interactions between biomolecules *in vitro* and *in vivo*. Improvements of fluorescent probes and labeling technologies in conjunction with instrumental improvements have enabled measurements that provide critical insights into cellular organization and biomolecular interactions occurring within cells^1^. Spatial co-localization of emitters through multi-color imaging has been widely utilized to gauge whether biomolecules are close in space and thus able to interact. The power of such measurements depends critically on the achievable spatial resolution of the instrument used. Diffraction-limited imaging provides 200-300 nm resolution, which is orders of magnitude larger than the size of typical proteins (~2 nm) or the size of large protein complexes (~20 nm). Diffraction-limited resolution is thus too low to conclusively determine whether two proteins interact directly or whether their interaction is mediated by a third protein^2^. Super-resolution microscopy approaches, such as PALM/STORM and MINFLUX, have been successful in addressing this challenge by enabling precise single-molecule localization with nanometer precision. For example, Symborska *et al*. determined the radial positions of protein subunits of the nuclear pore complex (NPC) with subnanometer precision using particle averaging of PALM/STORM images^3^. This approach was later extended to obtain a 3D reconstruction of the nuclear pore complex using iterative multi-color imaging of each NPC subunit relative to a reference protein within the NPC^4^. More recently, Ries and co-workers used MINFLUX microscopy to pinpoint the position of subunits within the NPC with single nanometer precision without the need for radial averaging^5^. Thus, for relatively large, immobile, and highly symmetric structures like the nuclear pore complex, super-resolution fluorescence microscopy can provide accurate co-localization data that provides information on how individual subunits interact with each other^6–8^.

Similar to the assembly of static complexes, biomolecular interactions in the cytosol can lead to freely diffusing complexes of varying molecular composition. Identifying and quantifying the relative abundances of fully and partially assembled cytosolic complexes in living cells can give valuable insights into the interaction networks that regulate biological processes *in vivo*^9,10^. For mobile complexes in living cells, spatial co-localization measurements using dual-color single-molecule localization microscopy are no longer practically feasible, because both fluorophores used to label the interacting proteins would have to be in a fluorescent ON state at the same time within the same complex. Fluorophore photo-activation or blinking in PALM/STORM occurs stochastically, such that the probability of observing co-localization simultaneously in separate color-channels is negligibly small. To avoid this problem, the diffusive behaviors of interacting proteins can be measured in separate single-molecule tracking experiments. If the two proteins diffuse at the same rate only when expressed simultaneously, then it is possible that they are part of the same complex. Because single-molecule trajectories in living cells are limited in duration by fluorophore photobleaching, information from thousands of trajectories are pooled and statistical data analysis approaches employed to resolve different diffusive states and determine their relative abundances and diffusion coefficients^11–17^. Using this approach, homo- and hetero-oligomeric complex formation among interacting proteins^18–21^ or proteins binding to quasi-stationary structures (such as DNA)^22–33^ have been successfully resolved in living cells.

Other fluorescence imaging-based methods, such as Förster Resonance Energy Transfer (FRET), Fluorescence Correlation Spectroscopy (FCS), and Fluorescence Recovery After Photobleaching (FRAP), can also be used to detect cytosolic protein interactions in living cells.

FRET efficiency serves as a population-averaged measurement of spatial proximity of fluorescently labeled proteins over a distance of a few nanometers^34,35^, and can thus be used to infer protein-protein interactions. While applied at the single-molecule level routinely *in vitro*, single-molecule FRET measurements in living cells require a series of complicated sample preparation protocols^36^. FRET measurements in living cells thus typically provide ensemble-averaged data that do not resolve individual diffusive states. Similarly, FRAP and FCS can be used to determine the average diffusion rate of proteins^37–39^, which is not amenable to quantifying the abundances of different protein complexes. Single-molecule tracking has thus emerged as the method of choice for obtaining diffusive state-resolved insights into cytosolic interaction networks that regulate biological processes in living cells.

Additional experimental perturbations are often needed to conclusively assign diffusive states to biomolecular complexes of specific protein composition. Single-point mutations or genetic deletion mutants of potential interacting partners can be used to disrupt complex formation and observe the resulting changes in diffusive behavior^11,12,14,16–19,21,23–33,40,41^. A drawback of such genetic approaches is that the permanent disruption of an interaction interface or absence of an essential binding partner can interfere with cellular processes in ways that are difficult to predict or control, and may compromise cell viability^42,43^. In such cases, experimentally controllable, transient perturbations of cellular processes at short time scales is preferable. Optically and chemically induced dimerization enables such transient perturbations and thus provides a promising alternative to permanent genetic approaches.

Optogenetic and chemogenetic dimerization systems enable manipulation of protein spatial localization to specific non-native cellular compartments in so-called knocksideways assays^44,45^. In a knocksideways experiment, one of the dimerizing molecules is attached to a cytosolic protein of interest and the other dimerization partner is targeted to a cellular location that differs from the native localization of the protein of interest (e.g. the cell membrane). Upon optical or chemical stimulation, dimerization is induced and the protein of interest, along with its interacting partners, is sequestered away from its native, cytosolic location. Chemogenetically induced dimerization relies on activation by a diffusing molecule, whereas optogenetic systems can be activated by photon absorption in real time^46^. Optogenetic systems thus allows for more precise spatial and temporal control over dimerization than chemogenetic systems.

When combined with single-molecule tracking, manipulating the spatial distributions of otherwise cytosolic proteins could aid the assignment of specific diffusive states to complexes of specific protein composition. In the simplest scenario, the diffusive state corresponding to a hetero-oligomer would be depleted when either interacting partner is sequestered to the membrane. Optogenetic knocksideways experiments can thus be used to directly detect interactions among cytosolic proteins in living cells.

Here, we test the possibility of combining optogenetic manipulation with 3D single-molecule tracking microscopy in live *Escherichia coli*. We selected the <u>i</u>mproved <u>L</u>ight <u>I</u>nduced <u>D</u>imerization (iLID) system^4^, because it has been extensively characterized and a large toolbox of iLID variants, each with different affinities, and reversion times, is available for researchers^47–49^. The iLID protein contains the light sensing light oxygen voltage (LOV2) domain derived from the oat cereal grass *Avena sativa*. The LOV2 domain incorporates a flavin cofactor during folding, which then acts as a chromophore and forms a cysteine adduct with the LOV2 domain after illumination with blue light. As a result, the iLID protein changes conformation and exposes a binding site for the interacting partner, SspB^50,51^. We demonstrate that the iLID system can be used to elicit a robust knock-sideways response after 488 nm illumination in low-intensity excitation conditions used for diffraction-limited imaging. Using single-molecule tracking of dye-labeled SspB, we found that SspB_micro_ exhibited the most substantial 488 nm-induced knock-sideways response among three tested SspB mutants. Surprisingly, our results also reveal that pre-activation of the optogenetic response occurs under high intensity laser illumination, even at wavelengths for which iLID shows minimal absorbance. Titrating the iLID expression level with respect to SspB reduced imaging laser-induced iLID:SspB interaction in 3D single-molecule tracking experiments. Together, these results establish the need for careful calibration of the optogenetic system prior to its application in quantitative imaging experiments for biological hypothesis testing.

## RESULTS AND DISCUSSION

### Diffraction-limited imaging indicates a robust redistribution of cytosolic proteins to the membrane upon blue-light activation of the iLID system

Optogenetic tools have been widely applied in eukaryotic cells in conjunction with diffraction-limited fluorescence microscopy to obtain population-averaged, phenotypic readouts^52,53^. Therefore, we sought to quantify the optogenetic response of the iLID system in K12 *E.coli* using diffraction-limited imaging as a first step. We expressed the iLID protein from an inducible arabinose promoter and targeted it to the inner membrane using an N-terminal genetic fusion containing a modified single transmembrane spanning helix derived from the *E.coli* transmembrane protein, TatA^54,55^. This construct, hereafter referred to as MA-iLID, was previously used for stable insertion into the inner membrane of Gram negative bacteria and has also been shown to enable optogenetic disruption of the type 3 secretion system (T3SS) in *Yersinia enterocolitica*^56^. We genetically fused the strongest reported iLID binding partner, SspB_nano_, with a C-terminal Halo tag, and expressed this construct from a second plasmid using a constitutively active promoter. SspB_nano_-Halo was labeled with JFX549 for fluorescence imaging^57^ (**Figure 1a**). In what follows, JFX549-labeled SspB_nano_-Halo is referred to as SspB_nano_ for simplicity.

**Figure 1.**
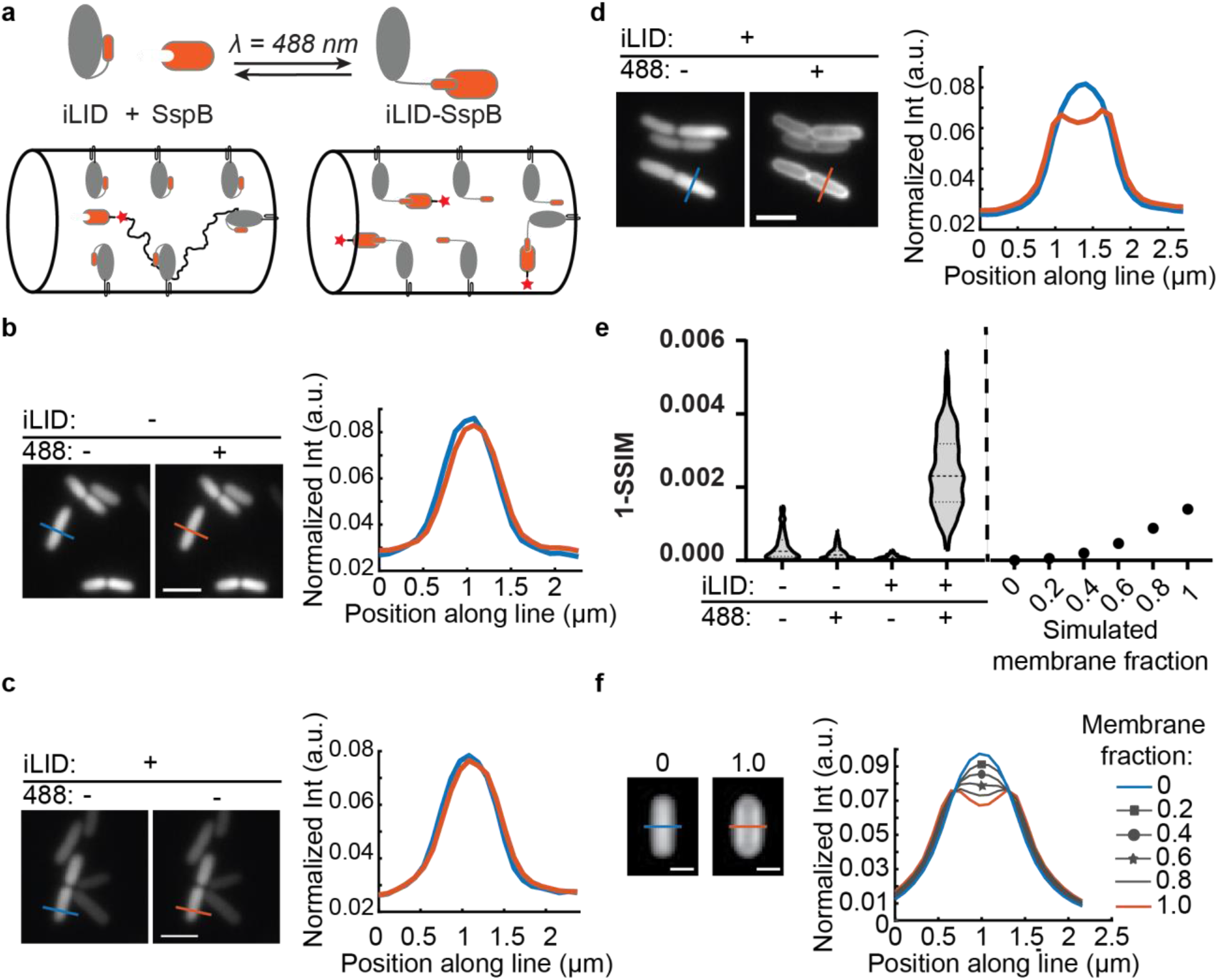
Diffraction-limited imaging indicates a substantial redistribution of SspB_nano_ to the membrane upon blue-light illumination. **(a)** Schematic depiction of a knocksideways experiment using the iLID system. The iLID protein undergoes a conformational change upon 488 nm-light illumination and uncages a binding site for SspB. Halo-tagged (red star) cytosolic SspB binds to the uncaged binding sites of membrane-anchored iLID (MA-iLID). SspB-Halo is labeled with JFX549 for fluorescence imaging. **(b)** Diffraction-limited images of SspB_nano_. Illumination with 488 nm light has no effect on the spatial distribution of SspB_nano_ in the absence of MA-iLID. Fluorescence intensity line profiles across the midsection of the cell (normalized to the integrated line intensity) also exhibit no apparent shift in SspB_nano_ spatial distribution. **(c)** In the presence of MA-iLID, low intensity (~1 W/cm^2^) 561 nm laser excitation over the entire exposure (80 seconds) is not sufficient to induce SspB_nano_ sequestration to the membrane. **(d)** In the presence of MA-iLID, low intensity (~4 mW/ cm^2^) 488 nm laser illumination sequesters SspB_nano_ to the cell membrane. The fluorescence intensity line profile across the midsection of the cell changes from a Gaussian-like line shape to a characteristic double-peaked line shape consistent with membrane association. **(e)** Left panel: Image dissimilarity (1-SSIM) of experimentally-acquired individual cell images before and after 488 nm illumination (*N*=200-350 cells for each condition). A pronounced population-level response in image dissimilarity requires both MA-iLID expression and 488 nm illumination. Right panel: Image dissimilarity metric obtained for simulated images with increasing fractions of membrane-associated fluorophores (relative to a simulated cell with 0% membrane-associated fluorophores). Variation in cell length and sample drift are not included in the simulated images, but contribute to the observed distribution of experimental dissimilarity values. **(f)** Simulated-diffraction-limited images of cells with 0% or 100% membrane-associated fluorophores, respectively. Normalized line profiles of simulated diffraction-limited images (**SI Figure 1**) indicate a progressive broadening, and redistribution of fluorescence as the fraction of membrane-associated fluorophores increases. Scale bars for images in **b-d** is 2.5 μm and 1 μm for simulated images in **f**.

Prior to optical activation of MA-iLID with 488 nm laser light, we observed a uniform distribution of cytosolic SspB_nano_. Spatial redistribution of SspB_nano_ to the membrane was dependent on both expression of MA-iLID and 488 nm laser illumination (~1mW/cm^2^) (**Figure 1b-d**). We quantified SspB_nano_ redistribution upon 488 nm laser illumination in each cell using the structural similarity index measure (SSIM), which quantifies similarity between two images^58^. Cells expressing MA-iLID were more dissimilar (1-SSIM) after 488 nm laser illumination compared to controls (**Figure 1e**). To estimate the fraction of molecules that reside at the membrane in each condition, we simulated 2D diffraction-limited images of cells with different fractions of membrane-associated fluorophores (see Methods) (**Figure 1f, SI Figure 1a**). Fluorescence line profiles across the midsection of cells showed a Gaussian-like line shape for cytosolic fluorescence, while membrane-associated fluorescence produced double-peaked line shapes. The experimental line shape obtained with a fluorescently labeled, membrane-anchored iLID protein (MA-mCherry-iLID) agrees well with line shape obtained based on simulated images of 100% membrane localized molecules (**SI Figure 1b**). Based on qualitative comparison of simulated and measured line shapes, we estimate that the fraction of molecules at the membrane in living cells is close to 0% pre-activation and greater than ~80% post-activation. We note that the degree of spatial redistribution is heterogeneous across the cell population (**SI Figure 2**). This may be due to expression level heterogeneity which is evident in whole cell fluorescence comparisons of both arabinose-promoter driven expression of MA-iLID, and constitutive expression of SspB (**Figure 1, SI Figure 2**). Variation in cell length and sub-pixel sample drift further add to the width of the experimentally-measured dissimilarity distribution – an effect that is not modeled in the simulated images (**Figure 1e** and **f**). Despite this heterogeneity, the obtained results establish that the iLID system can be used to efficiently sequester otherwise cytosolic molecules to the membrane in a knocksideways experiment.

### 3D single-molecule tracking data show increased SspB:iLID interaction prior to 488 nm illumination

Because blue-light activation of the MA-iLID:SspB interaction allows for spatial redistribution of cytosolic proteins, we reasoned that the iLID system could provide a means to transiently deplete specific diffusive states that manifest in the cytosol of living cells^59^. We therefore used 3D single-molecule tracking microscopy to measure the diffusive behavior of SspB in living cells and determine the extent of MA-iLID association. Based on the clear membrane localization of MA-iLID in diffraction-limited images (**Figure 1de** and **SI Figure 1**), we reasoned that SspB_nano_ interacting with MA-iLID should diffuse along the membrane at a much slower rate than cytosolic SspB_nano_. In 488 nm illuminated cells, the cumulative distribution function of apparent diffusion coefficients indeed shows a bimodal curve with a transition at *D*\* ~ 0.15 μm^2^/s (**Figure 2a**). The spatial trajectories of slowly diffusing molecules (*D*\* < 0.15 μm^2^/s) clearly localize near the bacterial membrane, while spatial trajectories of fast diffusing molecules (*D*\* > 0.15 um^2^/s) localize in the cytosol (**Figure 2b**). These result show that it is possible to clearly distinguish freely-diffusing, cytosolic SspB_nano_ from iLID-associated SspB_nano_ molecules in living cells based on their diffusion rate and sub-cellular localization.

**Figure 2.**
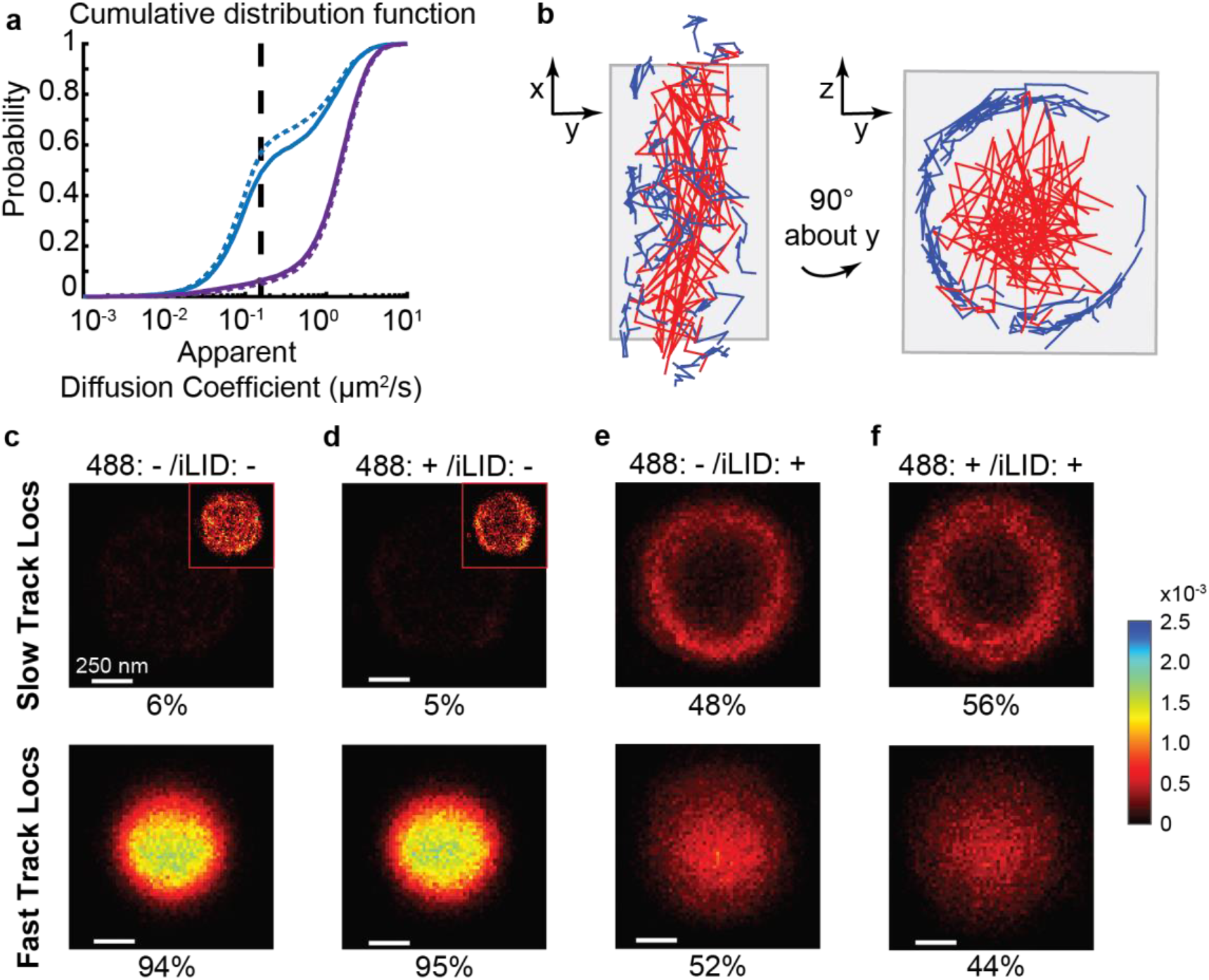
3D single-molecule tracking data show SspB:iLID interaction prior to 488 nm illumination. **(a)** Apparent diffusion coefficient distributions of SspB_nano_ in the absence (purple, solid line, *N* = 11,523 tracks, 120 cells) and presence (blue, solid line, *N* = 8,542 tracks, 78 cells) of MA-iLID. Illumination with 488 nm light marginally increases the population of slowly diffusing SspB_nano_ in MA-iLID expressing cells (blue, dashed line, *N* = 6,360 tracks, 88 cells), but not in cells which do not express MA-iLID (purple, dashed line, *N* = 9,922 tracks, 94 cells). Vertical dashed line indicates threshold used to distinguish fast diffusers (*D*\* > 0.15 μm^2^/s) from slow diffusers (*D*\* < 0.15 μm^2^/s) **(b)** Single-molecule trajectories in a representative cell expressing MA-iLID, after illumination with 488 nm light. Left: top view. Right: end-on view. Slowly diffusing molecules (blue) clearly localize to the periphery of the cell and undergo 2D diffusion near the membrane, while fast diffusing molecules (red) localize predominately to the cytosol and undergo 3D diffusion. **(c-f)** 2D histograms of localizations from slow and fast trajectories for the entire population of imaged cells. Histograms recapitulate trends observed in (b). Bin size 20 nm × 20 nm. Each histogram is normalized to the total number of molecules observed in that experiment. Inset: rescaled (unnormalized) histograms, shown for clarity.

The *D*\* = 0.15 um^2^/s threshold allowed us to estimate the relative population fractions of iLID-associated and cytosolic SspB_nano_. In cells that do not express MA-iLID, the vast majority (95%) of SspB_nano_ proteins diffuse fast and localize to the bacterial cytosol, as expected. The remaining 5% of tracked proteins were classified as slowly diffusing, but the corresponding trajectories showed no discernable preference for the membrane (**Figure 2c** and **d**). The diffusive behavior and spatial distribution of SspB_nano_ remained unchanged upon 488 nm illumination in these cells (**Figure 2a** and **d**). Surprisingly, and in stark contrast to the diffraction-limited images (**Figure 1**), a large fraction of SspB_nano_ proteins exhibited slow diffusion (48%) in cells that express MA-iLID, even in the absence of 488 nm light (**Figure 2a**). This shift towards slow diffusion is accompanied by spatial redistribution of SspB_nano_ to the membrane, whereas the fast diffusing population of proteins remained uniformly localized to the cytosol (**Figure 2e**). When cells expressing both SspB_nano_ and MA-iLID were exposed to 488 nm light, we observed an additional shift towards slower diffusion and membrane-proximal localization (56%) (**Figure 2a** and **f**). The increase in iLID-associated SspB observed using single-molecule tracking (**Figure 2**) is marginal compared to the shift observed in diffraction-limited images (**Figure 1**).

We hypothesize that the discrepancy in the fraction of MA-iLID associated SspB seen in diffraction-limited images and single-molecule trajectories is due to the higher excitation intensity required for single-molecule localization microscopy. The absorption spectrum of iLID proteins shows minimal absorption at wavelengths larger than 500 nm^60^. However, optical activation of the iLID:SspB_nano_ interaction is possible with 514 nm light at illumination intensities typical for diffraction-limited imaging (~1 W/cm^2^, **SI Figure 3**). The intensity used for single-molecule imaging of JFX549 using 561 nm laser light was three orders of magnitude higher (~2 kW/cm^2^). The results obtained here show that the increased 561 nm photon flux activates the iLID optogenetic response.

### SspB mutants exhibit differing degrees of pre-activation and optogenetic responses

To decrease the amount of iLID-bound SspB prior to 488 nm illumination, we made a series of previously characterized mutations to SspB^47,48^. These mutant versions of SspB, termed SspB_micro_ and SspB_milli_, have decreased binding affinity for the iLID protein. When we tracked single SspB_micro_ and SspB_milli_ proteins in living cells, we indeed observed a decrease in the fraction of slow diffusers near the cell membrane prior to 488 nm light illumination (**Figure 3a-c**). We found that 31% of SspB_micro_ proteins were iLID-associated, while iLID-associated SspB_milli_ comprised only 6% of the total population. Upon 488 nm light illumination, the membrane-proximal, slow-diffusing fractions increased to 45% and 7%, respectively (**Figure 3b** and **c**). The magnitude of the optogenetic response thus differs between the mutants, with SspB_micro_ showing the largest response, under otherwise identical experimental conditions. Diffraction-limited images of SspB_micro_ and SspB_milli_ recapitulate this trend (**SI Figure 4**). Previous work also shows that the SspB_milli_:iLID interaction is only minimally affected by blue light, as judged based on SspB_milli_ localization to iLID-labeled organelles in HeLa cells^61^.

**Figure 3.**
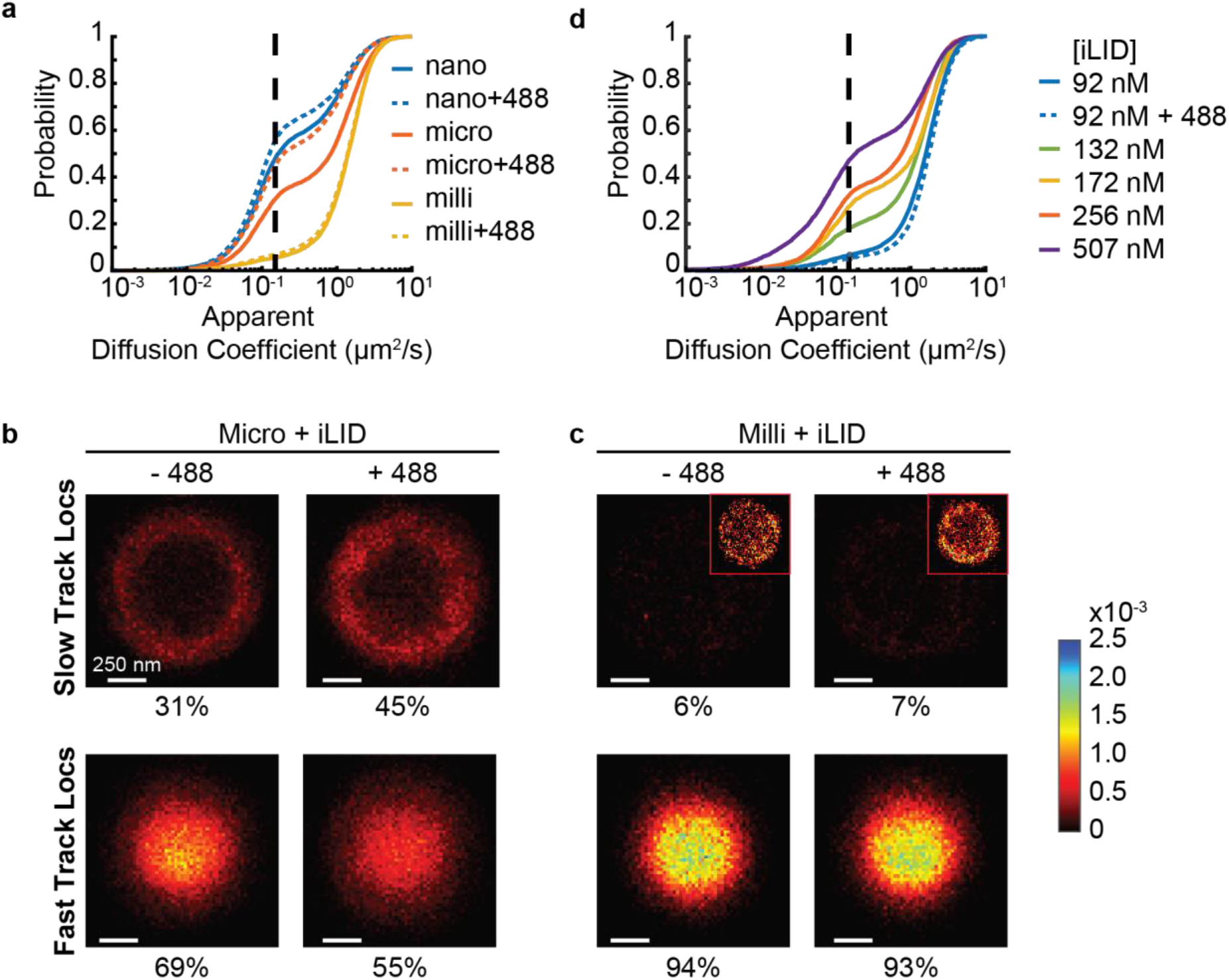
Affinity of the SspB:iLID interaction determines the fraction of iLID-associated SspB observed in single-molecule tracking experiments. **(a)** Apparent diffusion coefficient distributions for three SspB mutants. The lowest affinity mutant, SspB_milli_ (solid, yellow line, *N* = 4246 tracks, 65 cells) displays the highest population fraction of fast, cytosolic diffusion prior to 488 nm light illumination, followed by SspB_micro_ (solid, orange line, *N* = 7,738 tracks, 97 cells) and SspB_nano_ (solid, blue line, *N* = 8,542 tracks, 78 cells). Activation of these systems with 488 nm light (dashed line distributions) shifts each mutant protein’s distribution towards larger fractions of slow diffusing molecules due to MA-iLID association. **(b)** 2D cross-sectional histograms of SspB_micro_ trajectories indicate MA-iLID association at the membrane increases upon 488 nm light illumination (*N* = 6,491 tracks, 77 cells). Each histogram is normalized to the total number of molecules observed in that experiment. Inset: rescaled (unnormalized) histograms, shown for clarity. **(c)** 2D cross-sectional histograms of SspB_milli_ trajectories indicates small, but qualitatively discernable MA-iLID association upon 488 nm light illumination (*N* = 5243 tracks, 99 cells). **(d)** Apparent diffusion coefficient distribution of SspB_micro_ as a function of MA-iLID expression level. Higher MA-iLID expression leads to increased population fractions of slow, membrane-proximal diffusion of SspB_micro_ (see also **SI Figure 5** and **7**).

The reported dissociation constants for the three abovementioned SspB mutants differ by six orders of magnitude. The data reported above was collected at the same expression levels for each interacting protein components. SspB_nano_ may thus saturate the available binding sites at the membrane at the expression level used. Consequently, a large increase in binding upon 488 nm illumination is not observed. At the other extreme, SspB_milli_ may require a much higher number of binding sites to observe binding both before and after 488 nm illumination due to its inherently low affinity for the iLID protein. The comparatively larger optogenetic response observed for SspB_micro_ suggests that the concentration of each interacting parter is better-matched to the dissociation constant for this mutant.

To test this model, we titrated the amount of MA-iLID, while keeping the expression level of SspB_micro_ constant. To estimate the concentration of MA-iLID expressed at each L-arabinose induction concentration, we performed a calibration experiment in which we expressed fluorescent MA-mCherry-iLID under the same promoter. The number of MA-mCherry-iLID proteins was estimated by dividing the intial fluorescence intensity of each cell by the average intensity of a single mCherry protein. Uninduced cells still expressed MA-mCherry-iLID at an estimated concentration of ~90 nM due to leaky expression from the arabinose promoter (**SI Figure 5**). At the highest inducer concentration (13.3 mM), MA-mCherry-iLID reached a concentration of ~510 nM. For the following analyses, we assume that MA-mCherry-iLID and MA-iLID are expressed at the same level for a given L-arabinose concentration.

Decreasing the concentration of MA-iLID, decreases the fraction of iLID-associated SspB_micro_. At the lowest MA-iLID concentration, SspB_micro_ does not show an appreciable population fraction of slowly diffusing, membrane-proximal molecules both before and after 488 nm light illumination (**Figure 3d**). At the highest concentration of MA-iLID, we estimate that 48% of SspB_micro_ is iLID-associated. It is possible that this fraction could be augmented by further increasing the expression level of MA-iLID. However, we found that high expression of MA-mCherry-iLID leads to an increase in fluorescent foci formation, most often at the cell pole (**SI Figure 6a**). Fluorescent foci formation upon protein overexpression can be indicative of protein aggregation and/or formation of inclusion bodies^62,63^. These processes do not necessarily interfere with protein folding and function^62,63^. Indeed, we observe fluorescent foci formation in the MA-mCherry-iLID data, but not to the same extent in the SspB_micro_-Halo data (**SI Figure 6b**). These observations suggest that mCherry remains correctly folded and functional within the MA-mCherry-iLID construct. However, the iLID protein within the fluorescent foci may not be functional or fully available for interactions with SspB. Indeed, SspB_micro_-Halo shows a decreased propensity for fluorescent foci formation compared to the fraction observed for MA-mCherry (**SI Figure 6c**). Due to increased levels of fluorescent foci formation, we did not explore higher expression levels of MA-iLID.

Titration of MA-iLID expression allowed us to estimate the dissociation constant *K_D_* of the SspB_micro_:MA-iLID interaction in living cells. Because we observe 48%association of SspB_micro_ at 507 nM MA-iLID, we estimate that the *K_D_* is approximately 510 nM before 488 nm illumination. This value is about 100× smaller than the *K_D_* reported in *in vitro* fluorescence polarization assays in unactivated conditions^47^. However, as we have shown above, the high intensity 561 nm excitation light activates the iLID optogentic response. Therefore, it may be more appropriate to compare results to the published fluorescence polarization data obtained under activated conditions. Indeed, the reported *in vitro K_D_* is about 800 nM^47^, which is within a factor of two of the *K_D_* value we estimate here in living cells. We observe ~45% association of SspB_micro_ at ~260 nM MA-iLID in activated conditions, which represents an approximately two-fold increase in affinity upon 488 nm illumination. By contrast, *in vitro* fluorescence polarization assays reported a 58-fold change in binding affinity upon optical-activation of the system. While the increased 561 nm photon flux is responsible for activating the optogenetic response and thus reducing the range of observed affinity constants upon 488 nm illumination, we note that proteins reconstituted in lipid bilayers have also been shown to exhibit a shortened dynamic range of binding affinity. In those experiments, only a 2.7-fold change in affinity upon iLID-activation was observed^64,65^. These results suggest that proximity to the membrane, and perhaps the steric hindrance imposed by the membrane, may also play a role in the dynamics of the SspB:iLID interaction. Additional experiments may be able to differentiate the magnitude of these two effects.

### Cumulative displacement analysis identifies transient state-switching events in full-length trajectories

Previous analyses were performed on trajectories which were shortened to contain at most 10 displacements to prevent averaging over different diffusive states. Because our data sets contained many long trajectories (>10 displacements), we reasoned that some of these trajectories may contain information about the length of time a single SspB protein remains bound to an MA-iLID protein at the membrane. To assess the diffusive behavior over the course of a trajectory, we first calculated the cumulative displacement of each single-molecule over time (**Figure 4a** and **4b**, see methods). A similar method has been utilized previously to distinguish dimerization rates of membrane-associated proteins^66^. These data show a clear separation between fast, cytosolic diffusion and slow, membrane-associated diffusion. To estimate the diffusion coefficients that describe motion in each state, we simulated cytosolic and membrane-bound diffusion at different diffusion rates. We found that the cumulative displacement over time of single-molecule trajectories simulated at 5.5 μm^2^/s in the cytosol and 0.2 μm^2^/s at the membrane best matched the experimentally measured slopes of the cumulative displacements over time. The slope magnitudes were then used as references values for subsequent analysis.

**Figure 4.**
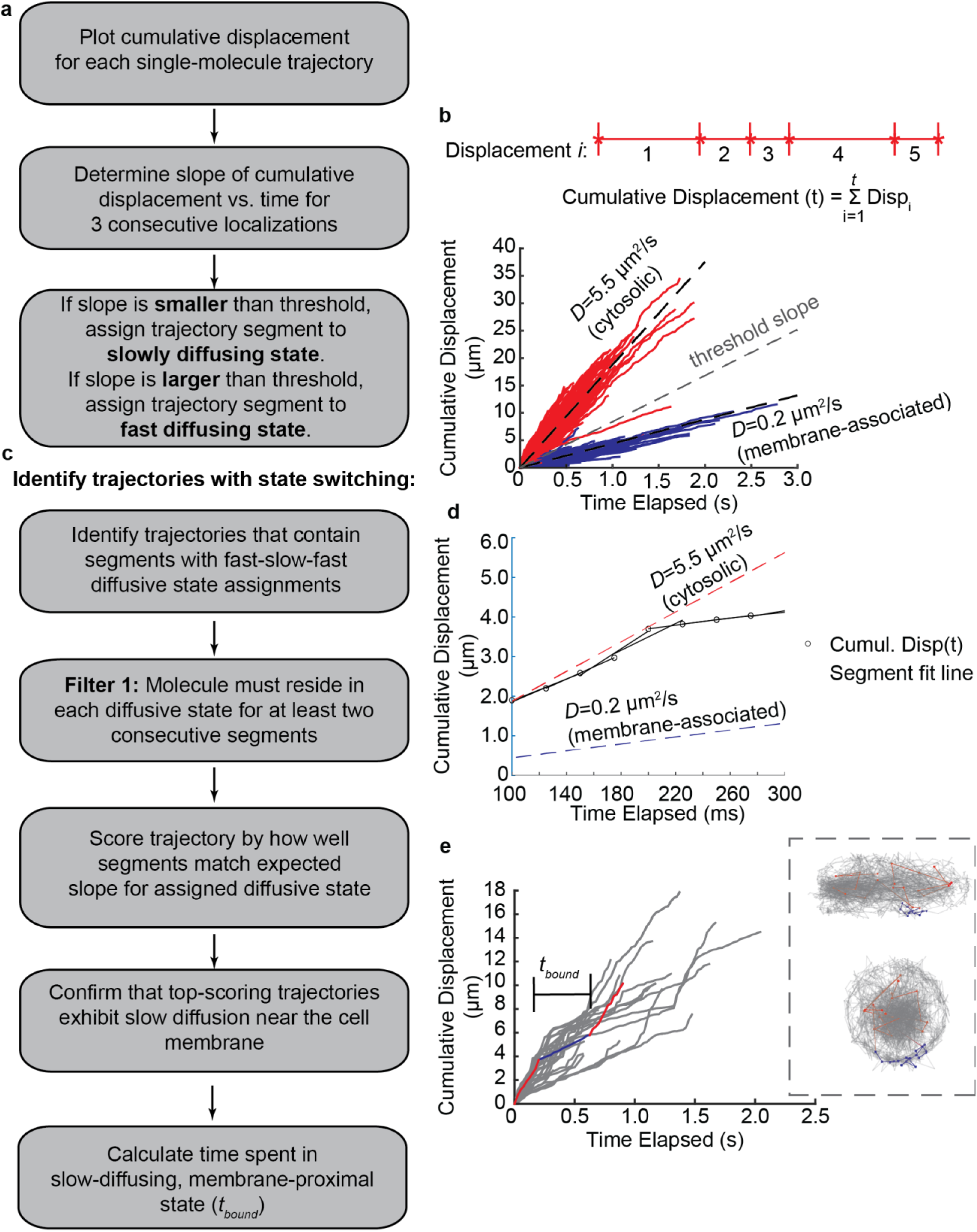
Cumulative displacement analysis identifies state-switching events in single-molecule trajectories. **(a)** Workflow of cumulative displacement analysis to distinguish fast, cytosolic diffusing molecules from slow, membrane-associated molecules. **(b)** Top panel: Schematic of displacement data transformed into a 1D trajectory to illustrate displacement size over time. The cumulative displacement is the sum of all displacements up to a given time point, *t*. Bottom panel: Cumulative displacement over time for SspB_micro_ trajectories collected in MA-iLID expressing cells (256 nM) in unactivated conditions. A threshold slope corresponding to *D* = 1.75 μm^2^/s was used to distinguish fast diffusing molecules (red) from slow diffusing molecules (blue). Simulated trajectories in the cytosol at 5.5 μm^2^/s and in the membrane at 0.2 μm^2^/s recapitulated experimental cumulative displacement slopes and were used as reference slopes in the following steps. **(c)** Workflow used to identify trajectories with state-switching (see text for details). **(d)** Example of a SspB_micro_ partial trajectory which exhibits state-switching. Three consecutive cumulative displacements are fit to a line, and the slope is compared to the threshold slope to classify the trajectory segment as cytosolic or membrane-bound. The slope of the segment is then compared to the reference slope for cytosolic diffusion or membrane diffusion depending on the state assignment, and scored by how closely they match. Trajectories with at least 2 consecutive segments in cytosolic and then membrane-associated states (or vice versa) are categorized as state-switching and used for later analysis. **(e)** Cumulative displacements over time of trajectories which show state transitions of fast-slow-fast. Inset: Example trajectory exhibiting fast(blue)-slow(red)-fast(blue) state-switching with the slowly diffusing trajectory segment near the membrane of the cell. Time spent in bound state (red portion of trajectory) determines the residence time *t_bound_*. **(f)** Histogram of residence times of SspB affinity mutants (bin size = 250 ms). The residence times seem to be independent of the affinity of the SspB:iLID interaction.

To identify state switching events within trajectories, we employed the workflow outlined in **Figure 4a** and **c**. Briefly, the cumulative displacements of small segments, containing 3 displacements, within trajectories were designated as fast diffusion or slow diffusion using a threshold corresponding to *D* = 1.75 μm^2^/s. This threshold was chosen to match the upper-limit of the displacement sizes used in the simulated, membrane-bound trajectories at 0.2 μm^2^/s. Experimental trajectories which exhibited subsequent fast-slow-fast diffusive states were ranked by evaluating the similarity of the experimental slope of cumulative displacements over time to the abovementioned reference slopes that described the simulated trajectories diffusing at *D* = 5.5 μm^2^/s and 0.2 μm^2^/s (**Figure 4d**). We next verified that the best scoring trajectories indeed exhibited slow-diffusing segments at the membrane, and fast-diffusing segments in the cytosol. The resulting subset of trajectories was then used to calculate the length of time, *t_bound_*, that a single SspB molecule remains in a slowly diffusing, membrane-proximal state (**Figure 4e**). As a control, we performed the same analysis for SspB trajectories in the absence of MA-iLID, and, as expected, did not identify clear state switching signatures.

In the presence of 256 nM MA-iLID, SspB_micro_ and SspB_nano_ exhibit similar bound times independent of 488 nm illumination (median = 300 and 350 ms, respectively) while SspB_milli_ displayed comparatively shorter binding times (median = 175 ms) (**Figure 4e**). Our measurements of *t_bound_* thus do not show a clear trend that correlates with each SspB mutant’s affinity for the iLID protein or with photoactivation of iLID. We note that trajectories that were identified as acceptable for residence time measurements represent less than 1% of the total number of trajectories. Cumulative displacement analysis also identified a much larger number of trajectories with slow diffusing molecules that stayed bound for the entire duration of the trajectory (**Figure 4b**). The fraction of slow-diffusing molecules identified using cumulative displacement analysis recapitulated the fraction derived from apparent cumulative distribution analyses. (**Figures 2** and **3, SI Figure 8**). The length of the trajectories of fully-bound molecules ranges from 0.3 seconds up to 3 seconds (longest observed trajectory – **Figure 4b**). The long lengths of these trajectories (which are terminated by fluorophore photobleaching) suggest that SspB_micro_ proteins can remain iLID-bound for much longer than the median time scales estimated for *t_bound_*. Previously published dissociation half-life estimates of SspB_nano_ and SspB_micro_ with iLID are on the order of minutes^64,65^. Our single-molecule trajectories are thus not long enough to capture both binding and unbinding events of stable SspB:iLID interactions. These results suggest that short time scale interactions that we capture are representative of a transient binding mode(s), which may not be reflective of stable SspB:iLID binding.

## CONCLUSIONS

A notable advantage of optogenetic systems, such as iLID, is that dimerization can be induced in real time using light. Optically-induced dimerization is thus much faster and spatially more controllable than chemically-induced dimerization, which is a diffusion-limited process. Even so, optically-induced dimerization systems have to be carefully characterized prior to their application in biological systems. Here, we provide a quantitative analysis of the iLID optogenetic system in live K12 *E. coli* cells using 2D diffraction-limited imaging and 3D single-molecule tracking. We show that the iLID system enables efficient optogenetic manipulation of protein spatial distributions when used in conjunction with diffraction-limited imaging. Single-molecule tracking data acquired using high intensity 561 nm laser excitation however leads to a substantial amount of SspB:iLID interaction even in the absence of optogenetic activation at 488 nm. These findings show that increased photon flux is able to activate the iLID optogenetic response, even at wavelengths greater than 500 nm where the LOV2 domain absorption spectra show negligible absorption^60^. This conclusion is consistent with our observation that even low intensity 514 nm light illumination is able to increase the SspB:iLID interaction. Other optogenetic systems, which utilize the LOV2 light sensing domain, have also reported 514 nm light induced activation^67^. Another possible explanation is that proximity to the membrane and imposed steric hindrance from membrane-tethering results in a shift in iLID conformational dynamics, resulting in a change in kinetics of the SspB:iLID interaction *in vivo*. Indeed, a smaller change in the measured dissociation constants has been seen in other model systems utilizing the iLID:SspB interaction where one of the binding partners is surface-immobilized^64,65^.

Our results highlight the importance of calibrating the effects of illumination conditions and cellular environment when the iLID system, or any optogenetic system, is used to modulate the spatial distribution of SspB-tagged proteins. Although pre-activation dampens the magnitude of the optogenetic response that can be observed, highly sampled distributions obtained from tens of thousands of single-molecule trajectories, may allow for the detection of statistically significant amounts of cytosolic diffusive states depletion. Additionally, titration of protein expression could be used to incrementally sequester ever larger amounts of proteins of interest. To avoid pre-activation altogether, far-red excitable fluorophores could be employed for single-molecule tracking. Based on the trends observed here we speculate that high intensity red and far-red light illumination will not activate the iLID system. Thus, single-molecule localization and tracking microscopy with red and far-red fluorophores would likely enable observations of large optogenetic responses, similar to those observed in diffraction-limited imaging.

## METHODS

### Bacterial strains and plasmids

*Escherichia coli* K12 (MG1655) strains were generated by introducing pACYC and pBAD vectors containing genes encoding the cytosolic prey protein, SspB, and the membrane-anchored bait protein, iLID, respectively. pACYC SspB_nano_-Halo was derived from pACYC SspB_nano_-mCherry (pFL109) which was a gift from Andreas Diepold. The construct was modified by removing mCherry from the plasmid via restriction digest and the *halo* coding sequence was ligated in its place, using XhoI and SalI cut sites. The plasmid containing the bait protein, pBAD FLAG-iLID (pFL108), also a gift from the Diepold lab, was transformed into bacteria without any further modification.

### Cell culture

*E. coli* cultures were inoculated from a freezer stock and grown overnight in LB media at 37° C and shaking at 225 rpm. Strains expressing from the pACYC and pBAD plasmids were grown in 30 μg/mL chloramphenicol and 100 μg/mL ampicillin, respectively. Overnight cultures were diluted to an OD_600_ of 0.05 in M9 minimal media and incubated at 37° C. Strains containing the pBAD plasmid were induced with 5.33 mM L-arabinose after 2 hours unless indicated otherwise. After an additional 1.5 hours of growth, 1.5 mL of cell culture was aliquoted and stained with 1.5 μL of 500 μM JFX-549 dye. The cell suspension mixture was incubated at 37° C for 30 minutes and then washed four times with M9 minimal media. Cell pellets were resuspended in a final volume of 10 μL for use in imaging experiments.

*E. coli* cells were transformed with plasmids via heat shock of chemically competent cells. Transformed cells were selected for by growing on LB agar plates containing chloramphenicol, ampicillin (100 μg/mL), or a combination of both for co-expressing strains. Transformants were screened by performing a Western blot for FLAG or Halo proteins in pBAD and pACYC-containing strains, respectively. Strains which showed a clear band in Western blot experiments were used to make freezer stocks containing 15% glycerol.

### Optical setup

Imaging of cells was performed on a home-built inverted fluorescence microscope as described previously^21^. An additional 488 nm laser line (Coherent Genesis) was used for activation of the iLID system. Emitted fluorescence within a spectral window of 570 nm – 700 nm was captured using an sCMOS camera (Hamamatsu, ORCA Flash 4.0 v2).

### Epifluorescence imaging

Stained cells were mounted on a #1.5 coverslips (VWR) and immobilized using 1.5% agarose pad prepared in M9 minimal media. Halo-conjugated JFX549 was excited using 561 nm laser light at ~1 W/cm^2^. Images were acquired at 5 frames/sec for 40 seconds pre- and post-activation of the iLID optogenetics system. The iLID optogenetic system was activated using 488 nm laser light at ~4 mW/cm^2^ continuously for 2 minutes prior to and during acquisition of post-activation images. Epifluorescence images shown in the figures represent averages of these image sequences. Brightfield images were acquired of the same field-of-view by illuminating cells with an LED and taking a single 25 ms exposure.

### SSIM analysis of Diffraction-limited images

Pre- and post-activation images were background subtracted. Outlines of cells were derived by using OUFTI^68^ on inverted color to create a binary mask such that each cell was compared to itself before and after activation. The built-in Matlab (The MathWorks, Inc, Natick, Massachusetts) function ssim() was applied to the masked images with equal weighting of the comparison metrics, luminance, structure, and contrast, to assess the degree of image similarity. Cells with abnormal morphology (based on the associated brightfield image) were excluded from the analysis. Multiple fields-of-view were acquired for each condition, and the experiments were replicated twice.

### Super-resolution fluorescence imaging

Fluorescent fiducial markers (Invitrogen) were added to cell suspensions and the cell suspension was then mounted on #1.5 glass coverslips. Cells were immobilized using solidified pads of 1.5% agarose in M9 minimal media. Halo-conjugated JFX-549 was excited into the blinking state by 561 nm laser light at ~2 kW/cm^2^ at the sample. Images were acquired at 40 frames/sec in the presence or absence optogenetic activating 488 nm light. Cells were activated using 488 nm laser light at ~4 mW/cm^2^ continuously for 2 minutes prior to and during acquisition of post-activation images.

### Data Processing

Raw data was processed in Matlab (The MathWorks, Inc, Natick, Massachusetts) using a modified version of easy-DHPSF software^21,69^. Fluorescent fiducial markers were used for sample drift correction. Single-molecule localizations were registered to phase contrast images using a two-step affine transformation in MATLAB as previously described^21^. Localizations outside of cells were discarded from further analysis using axial bounds and OUFTI-derived cell outlines which were generated from phase contrast images, as described above.

### Single-molecule tracking analysis

To derive single-molecule displacements, localizations in subsequent frames were linked into trajectories. A maximum linking distance of 2.5 um was used for linking analysis, and multiple localizations which were present at the same time within a single cell were discarded to prevent misassignment of molecules.

The Mean Squared Displacement (MSD) for each trajectory was calculated using

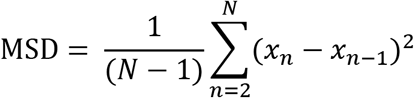

where *x* is the 3D position at timepoint *n*, including up to 11 timepoints for calculating the mean over 10 displacements. The remainder of the trajectory was not used in the MSD analysis to ensure multiple diffusive states were not averaged together. The MSD measurement was then used to calculate the apparent diffusion coefficient (*D*\*) according to:

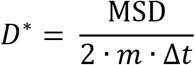

where *m* is the dimensionality of the measurement (*m*=3 for the 3D trajectories reported here), and Δ*t* is the camera exposure time used for imaging (Δ*t*=25 ms under our conditions).

### 2D-cross section projection analysis

The vector describing the central axis of each cell was determined using the cell outline generated from OUFTI. The outline was segmented into sections along the cell length, and localizations from trajectories in each section were projected onto a 2D plane. The position of the central cell axis was adjusted to match the centroid of all localizations within the section. Positions of localizations from each cell were scaled to match the mean cell radius and mean cell length which was calculated from OUFTI outlines. The trajectories were classified as slow or fast diffusing using the threshold *D*\* = 0.15 μm^2^/s, which shows the clearest separation between membrane-associated and cytosolic trajectories, as determined by visual inspection of trajectory localization (**Figure 2**).

### Determination of *t_bound_*

To identify diffusive state transitions in single-molecule trajectories, we developed a workflow to analyze displacement data over time (**Figure 4a and c**). First, we plotted the cumulative displacement (CD) as a function of time elapsed according to:

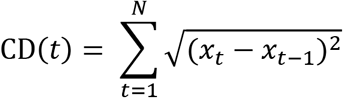

where *x* is the 3D position at timepoint *t*. The CD was calculated for each displacement, for the full trajectory length (*N* displacements).

The instantaneous rate of displacement was estimated by finding the slope within a small, sliding window containing 3 displacements for the full length of the trajectory (**Figure 4d**). The size of the sliding window was chosen to be small enough such that short duration state switching events were not excessively averaged, while being large enough to be somewhat insensitive to displacement size fluctuations due to localization error. Each segment of the trajectory was then classified as bound or free by calculating its slope and comparing it to a threshold, which was chosen to match the upper-limit of the displacement sizes used in the membrane-bound simulation at 0.2 μm^2^/s. Trajectories which contained consecutive segments of fast-slow-fast state assignments were used for further analysis. To be considered for analysis, the molecule must reside in each state for at least 2 segments (4 displacements) of the trajectory to ensure that the identified state change was a true diffusive state change and not an artefact due to localization error. Each switching trajectory was ascribed a score (*S*) quantifying how well the slope of trajectory segments matched simulated cumulative diffusion coefficient slopes derived for freely diffusing, cytosolic and membrane-bound molecules:

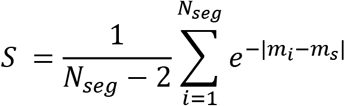

where *m_i_* is the slope fit to the experimental trajectory segment, and *m_s_* is the slope derived from free cytosolic or membrane-bound simulated data, depending on the state assigned to the trajectory segment. *N_seg_* refers to the number of segments in the trajectory. Switching trajectories were ranked by their TS score, and the top-scoring trajectories (TS>~0.16) were inspected individually to verify that switching events occurred at the membrane of the cell. The residence time (*t_bound_*) of bound molecules in these switching trajectories was calculated by determine the number of displacements the molecule was assigned to the bound state.

### 3D single-molecule trajectory simulation

Cytosolic trajectories were simulated as described previously.^19,21^ Briefly, Monte Carlo simulations of Brownian motion at 5.5 μm^2^/s were confined to the volume of a cylinder matching the dimensions of K12 *E. coli* (radius = 0.5 μm, length = 2.5 μm). Initial positions of molecules were randomly selected to uniformly fill the volume of the cell, and subsequent positions in the trajectory were selected at 100 ns intervals for a total of 125 ms. We simulated noisy, motion-blurred images of the single-molecule trajectories, integrating images over subsequent 25 ms time intervals, to match experimental camera exposure times for a total of 6 images per single-molecule track. We simulated 1000 such trajectories. To best match data derived from experimental conditions, we analyzed the simulated images of trajectories in the same way described for experimental localization and tracking. Easy-DHPSF software was used to analyze the images and detect single-molecule localizations.

Membrane-bound trajectories were simulated for molecules diffusing at 0.2 μm^2^/s. Initial positions were randomly chosen on the surface of a spherocylinder matching the dimensions of K12 *E. coli* (radius = 0.5 μm, length = 2.5 μm). Monte Carlo simulations of molecular motion were confined to the surface of the spherocylinder by translating displacements to changes in lateral position and elevation angle on the cylindrical surface or elevation and azimuthal angle on the hemispherical surface, such that the distance from the cell axis is always equal to the radius of the cell. Positions were updated at 100 ns intervals for a total of 125 ms. Images were generated and analyzed as described in the previous paragraph.

### 2D diffraction-limited image simulation

Trajectories of cytosolic and membrane-associated diffusing molecules were simulated as described above with the following modifications to match diffraction-limited imaging conditions. Subsequent positions of molecules were sampled every 1 μs for a total of 1.2 seconds to generate a finely sampled trajectory for molecules diffusing at 5.5 μm^2^/s in the cytosol and 0.2 μm^2^/s in the membrane. The positions of the molecule at 200 ms time intervals were convolved with diffraction-limited point spread functions of the microscope using a vectorial light propagation model^70^. A total of 10,000 trajectories were simulated for both cytosolic and membrane-bound molecules, providing 60,000 total emitter positions in each condition. We modified the algorithm to omit the double helix phase mask and optical aberrations and thus generate an ideal diffraction-limited image. In the simulation, the nominal focal plane was positioned at the center of the cell.

Images obtained for each single molecule were added to attain the total image. To attain mixed population images, trajectories were chosen at random from the total simulated trajectory population to reflect the specified membrane-associated molecule fraction.

## AUTHOR CONTRIBUTIONS

A. A., and A. G. designed research;

A. A. performed research;

A. A., T. Y. and A. G. analyzed data;

A. A. and A. G. wrote the paper.

## COMPETING INTERESTS

A. G. declares the following competing interests: A European patent application and PCT registration on the presented method was filed by the Max-Planck-Gesellschaft zur Förderung der Wissenschaften e.V.: Lindner, F., Gahlmann, A., Diepold, A. “Optogenetic control of protein translocation—Protein secretion and translocation into eukaryotic cells with high spatial and temporal resolution by light-controlled activation of the bacterial type III secretion system”, European Patent Application 19166308, March 2019 and PCT registration, March 2020. The remaining authors declare no competing interests.

## ACKNOWLEDGEMENTS

A. A. was supported in part by the U.S. National Institutes of Health Training Grant 5T32GM080186–08. We thank Andreas Diepold for providing parent bacterial strains and plasmids used in this work.

## Notes

### Summary of Updates

Main text updated to improve clarity. Added SI Figure 2. Acknowledgements, author contributions, and competing interests statements.

## REFERENCES

1 Achimovich, A. M., Ai, H. & Gahlmann, A. Enabling technologies in super-resolution fluorescence microscopy: reporters, labeling, and methods of measurement. Current Opinion in Structural Biology 58, 224–232, doi:https://doi.org/10.1016/j.sbi.2019.05.001 (2019).

2 Aaron, J. S., Taylor, A. B. & Chew, T.-L. Image co-localization – co-occurrence versus correlation. Journal of Cell Science 131, jcs211847, doi:10.1242/jcs.211847 (2018).

3 Szymborska, A. et al. Nuclear Pore Scaffold Structure Analyzed by Super-Resolution Microscopy and Particle Averaging. Science 341, 655–658, doi:10.1126/science.1240672 (2013).

4 Sieben, C., Banterle, N., Douglass, K. M., Gönczy, P. & Manley, S. Multicolor single-particle reconstruction of protein complexes. Nature Methods 15, 777–780, doi:10.1038/s41592-018-0140-x (2018).

5 Thevathasan, J. V. et al. Nuclear pores as versatile reference standards for quantitative superresolution microscopy. Nature Methods 16, 1045–1053, doi:10.1038/s41592-019-0574-9 (2019).

6 Malkusch, S. et al. Coordinate-based colocalization analysis of single-molecule localization microscopy data. Histochemistry and cell biology, doi:10.1007/s00418-011-0880-5 (2011).

7 Larson, J. D., Rodgers, M. L. & Hoskins, A. A. Visualizing cellular machines with colocalization single molecule microscopy. Chem. Soc. Rev. 43, 1189–1200, doi:10.1039/c3cs60208g (2014).

8 Levet, F. et al. A tessellation-based colocalization analysis approach for single-molecule localization microscopy. Nature communications 10, doi:10.1038/s41467-019-10007-4 (2019).

9 Szklarczyk, D. et al. The STRING database in 2021: customizable protein–protein networks, and functional characterization of user-uploaded gene/measurement sets. Nucleic Acids Research 49, D605–D612, doi:10.1093/nar/gkaa1074 (2021).

10 Fionda, V. in Encyclopedia of Bioinformatics and Computational Biology (eds Shoba Ranganathan, Michael Gribskov, Kenta Nakai, & Christian Schönbach) 915–921 (Academic Press, 2019).

11 Elf, J. & Barkefors, I. Single-Molecule Kinetics in Living Cells. Annual Review of Biochemistry 88, 635–659, doi:10.1146/annurev-biochem-013118-110801 (2019).

12 Karslake, J. D. et al. SMAUG: Analyzing single-molecule tracks with nonparametric Bayesian statistics. Methods 193, 16–26, doi:10.1016/j.ymeth.2020.03.008 (2021).

13 Hansen, A. S. et al. Robust model-based analysis of single-particle tracking experiments with Spot-On. eLife 7, 7 (2018).

14 Persson, F., Lindén, M., Unoson, C. & Elf, J. Extracting intracellular diffusive states and transition rates from single-molecule tracking data. Nature Methods 10, 265–269, doi:10.1038/nmeth.2367 (2013).

15 Monnier, N. et al. Inferring transient particle transport dynamics in live cells. Nature Methods 12, 838–840, doi:10.1038/nmeth.3483 (2015).

16 Chen, T. Y. et al. Quantifying Multistate Cytoplasmic Molecular Diffusion in Bacterial Cells via Inverse Transform of Confined Displacement Distribution. The journal of physical chemistry. B 119, 14451–14459, doi:10.1021/acs.jpcb.5b08654 (2015).

17 Michalet, X. & Berglund, A. J. Optimal diffusion coefficient estimation in single-particle tracking. Physical Review E 85, doi:10.1103/PhysRevE.85.061916 (2012).

18 Prindle, J. R., Rocha, J., Wang, Y., Diepold, A. & Gahlmann, A. Distinct complexes containing the cytosolic type III secretion system ATPase resolved by 3D single-molecule tracking in live <em>Yersinia enterocolitica</em>. bioRxiv, 2022.2004.2025.488798, doi:10.1101/2022.04.25.488798 (2022).

19 Rocha, J., Corbitt, J., Yan, T., Richardson, C. & Gahlmann, A. Resolving Cytosolic Diffusive States in Bacteria by Single-Molecule Tracking. Biophysical journal 116, 1970–1983, doi:https://doi.org/10.1016/j.bpj.2019.03.039 (2019).

20 Rocha, J. M. Dynamic Assembly of the Type-3 Secretion System in Yersinia enterocolitica Probed by Super-Resolution Fluorescence Imaging Doctor of Philosophy thesis, Graduate School of Arts and Sciences, University of Virginia, (2021).

21 Rocha, J. M. et al. Single-molecule tracking in live Yersinia enterocolitica reveals distinct cytosolic complexes of injectisome subunits. Integrative Biology 10, 502–515, doi:10.1039/c8ib00075a (2018).

22 Elf, J., Li, G. W. & Xie, X. S. Probing Transcription Factor Dynamics at the Single-Molecule Level in a Living Cell. Science 316, 1191–1194, doi:10.1126/science.1141967 (2007).

23 Gahlmann, A. & Moerner, W. E. Exploring bacterial cell biology with single-molecule tracking and super-resolution imaging. Nature Reviews Microbiology 12, 9–22, doi:10.1038/nrmicro3154 (2014).

24 Uphoff, S., Sherratt, D. J. & Kapanidis, A. N. Visualizing Protein-DNA Interactions in Live Bacterial Cells Using Photoactivated Single-molecule Tracking. JoVE, e51177, doi:doi:10.3791/51177 (2014).

25 Badrinarayanan, A., Reyes-Lamothe, R., Uphoff, S., Leake, M. C. & Sherratt, D. J. *In Vivo* Architecture and Action of Bacterial Structural Maintenance of Chromosome Proteins. Science 338, 528–531, doi:10.1126/science.1227126 (2012).

26 Stracy, M. et al. Single-molecule imaging of UvrA and UvrB recruitment to DNA lesions in living Escherichia coli. Nature communications 7, 12568, doi:10.1038/ncomms12568 (2016).

27 Uphoff, S. et al. Stochastic activation of a DNA damage response causes cell-to-cell mutation rate variation. Science 351, 1094–1097, doi:10.1126/science.aac9786 (2016).

28 Uphoff, S., Reyes-Lamothe, R., Garza de Leon, F., Sherratt, D. J. & Kapanidis, A. N. Single-molecule DNA repair in live bacteria. Proceedings of the National Academy of Sciences of the United States of America 110, 8063–8068, doi:10.1073/pnas.1301804110 (2013).

29 Liao, Y., Li, Y., Schroeder, J. W., Simmons, L. A. & Biteen, J. S. Single-Molecule DNA Polymerase Dynamics at a Bacterial Replisome in Live Cells. Biophysical journal 111, 2562–2569, doi:10.1016/j.bpj.2016.11.006 (2016).

30 Liao, Y., Schroeder, J. W., Gao, B., Simmons, L. A. & Biteen, J. S. Single-molecule motions and interactions in live cells reveal target search dynamics in mismatch repair. Proc Natl Acad Sci U S A 112, E6898–6906, doi:10.1073/pnas.1507386112 (2015).

31 Calkins, A. L. et al. Independent Promoter Recognition by TcpP Precedes Cooperative Promoter Activation by TcpP and ToxR. mBio 12, e0221321, doi:10.1128/mBio.02213-21 (2021).

32 Li, Y., Chen, Z., Matthews, L. A., Simmons, L. A. & Biteen, J. S. Dynamic Exchange of Two Essential DNA Polymerases during Replication and after Fork Arrest. Biophysical journal 116, 684–693, doi:10.1016/j.bpj.2019.01.008 (2019).

33 Li, Y., Schroeder, J. W., Simmons, L. A. & Biteen, J. S. Visualizing bacterial DNA replication and repair with molecular resolution. Curr Opin Microbiol 43, 38–45, doi:10.1016/j.mib.2017.11.009 (2017).

34 Zeug, A., Woehler, A., Neher, E. & Evgeni. Quantitative Intensity-Based FRET Approaches—A Comparative Snapshot. Biophysical journal 103, 1821–1827, doi:10.1016/j.bpj.2012.09.031 (2012).

35 Forster, T. Energiewanderung und fluoreszenz. Naturwissenschaften 33, 166–175 (1946).

36 Konig, I. et al. Single-molecule spectroscopy of protein conformational dynamics in live eukaryotic cells. Nat Methods 12, 773–779, doi:10.1038/nmeth.3475 (2015).

37 Magde, D., Elson, E. & Webb, W. W. Thermodynamic Fluctuations in a Reacting System—Measurement by Fluorescence Correlation Spectroscopy. Phys. Rev. Lett. 29, 705–708, doi:10.1103/physrevlett.29.705 (1972).

38 Axelrod, D. K., D. E.; Schlessinger J.; Elson, E.; Webb, W.W. Mobility measurement by analysis of fluorescence photobleaching recovery kinetics. Biophysical journal 16, 1055–1069, doi:https://doi.org/10.1016/S0006-3495(76)85755-4 (1976).

39 Koppel, D. E. A., D.; Schlessinger, J.; Elson, E. L.; Webb, W. W. . Dynamics of fluorescence marker concentration as a probe of mobility. Biophysical journal 16, 1315–1329 (1976).

40 Hansen, A. S. et al. Robust model-based analysis of single-particle tracking experiments with Spot-On. Elife 7, doi:10.7554/eLife.33125 (2018).

41 Monnier, N. et al. Inferring transient particle transport dynamics in live cells. Nat Methods 12, 838–840, doi:10.1038/nmeth.3483 (2015).

42 Peters, J. M. et al. A Comprehensive, CRISPR-based Functional Analysis of Essential Genes in Bacteria. Cell 165, 1493–1506, doi:10.1016/j.cell.2016.05.003 (2016).

43 Ye, Y.-N., Hua, Z.-G., Huang, J., Rao, N. & Guo, F.-B. CEG: a database of essential gene clusters. BMC Genomics 14, 769, doi:10.1186/1471-2164-14-769 (2013).

44 Robinson, M. S. & Hirst, J. Rapid inactivation of proteins by knocksideways. Curr Protoc Cell Biol 61, 15 20 11–17, doi:10.1002/0471143030.cb1520s61 (2013).

45 Benedetti, L. et al. Light-activated protein interaction with high spatial subcellular confinement. Proc Natl Acad Sci U S A 115, E2238–E2245, doi:10.1073/pnas.1713845115 (2018).

46 Kottke, T., Xie, A., Larsen, D. S. & Hoff, W. D. Photoreceptors Take Charge: Emerging Principles for Light Sensing. Annual Review of Biophysics 47, 291–313, doi:10.1146/annurev-biophys-070317-033047 (2018).

47 Guntas, G. et al. Engineering an improved light-induced dimer (iLID) for controlling the localization and activity of signaling proteins. Proc Natl Acad Sci U S A 112, 112–117, doi:10.1073/pnas.1417910112 (2015).

48 Zimmerman, S. P. et al. Tuning the Binding Affinities and Reversion Kinetics of a Light Inducible Dimer Allows Control of Transmembrane Protein Localization. Biochemistry 55, 5264–5271, doi:10.1021/acs.biochem.6b00529 (2016).

49 Lindner, F., Gahlmann, A. & Diepold, A. Optogenetic control of protein translocation: Protein secretion and translocation into eukaryotic cells with high spatial and temporal resolution by light-controlled activation of the bacterial type III secretion system. European Patent Application 19166308 patent (2019).

50 Harper, S. M. Structural Basis of a Phototropin Light Switch. Science (American Association for the Advancement of Science) 301, 1541–1544.

51 Zayner, J. P., Antoniou, C. & Sosnick, T. R. The Amino-Terminal Helix Modulates Light-Activated Conformational Changes in AsLOV2. J. Mol. Biol. 419, 61–74, doi:10.1016/j.jmb.2012.02.037 (2012).

52 Kramer, M. M., Lataster, L., Weber, W. & Radziwill, G. Optogenetic Approaches for the Spatiotemporal Control of Signal Transduction Pathways. International Journal of Molecular Sciences 22, 5300, doi:10.3390/ijms22105300 (2021).

53 Wittmann, T., Dema, A. & Van Haren, J. Lights, cytoskeleton, action: Optogenetic control of cell dynamics. Current Opinion in Cell Biology 66, 1–10, doi:10.1016/j.ceb.2020.03.003 (2020).

54 De Geyter, J., Smets, D., Karamanou, S. & Economou, A. 337–366 (Springer International Publishing, 2019).

55 Cline, K. Mechanistic Aspects of Folded Protein Transport by the Twin Arginine Translocase (Tat). Journal of Biological Chemistry 290, 16530–16538, doi:10.1074/jbc.r114.626820 (2015).

56 Lindner, F., Milne-Davies, B., Langenfeld, K., Stiewe, T. & Diepold, A. LITESEC-T3SS - Light-controlled protein delivery into eukaryotic cells with high spatial and temporal resolution. Nature communications 11, 2381, doi:10.1038/s41467-020-16169-w (2020).

57 Grimm, J. B. et al. A General Method to Improve Fluorophores Using Deuterated Auxochromes. JACS Au 1, 690–696, doi:10.1021/jacsau.1c00006 (2021).

58 Wang, Z., Simoncelli, E. P. & Bovik, A. C. (IEEE, 2003).

59 Elf, J. & Barkefors, I. Single-Molecule Kinetics in Living Cells. Annu Rev Biochem 88, 635–659, doi:10.1146/annurev-biochem-013118-110801 (2019).

60 Swartz, T. E. et al. The Photocycle of a Flavin-binding Domain of the Blue Light Photoreceptor Phototropin. Journal of Biological Chemistry 276, 36493–36500, doi:10.1074/jbc.m103114200 (2001).

61 Nijenhuis, W., Van Grinsven, M. M. P. & Kapitein, L. C. An optimized toolbox for the optogenetic control of intracellular transport. J. Cell Biol. 219, doi:10.1083/jcb.201907149 (2020).

62 García-Fruitós, E. et al. Aggregation as bacterial inclusion bodies does not imply inactivation of enzymes and fluorescent proteins. Microbial Cell Factories 4, 27, doi:10.1186/1475-2859-4-27 (2005).

63 Rinas, U. et al. Bacterial Inclusion Bodies: Discovering Their Better Half. Trends in Biochemical Sciences 42, 726–737, doi:10.1016/j.tibs.2017.01.005 (2017).

64 Bartelt, S. M. et al. Dynamic blue light-switchable protein patterns on giant unilamellar vesicles. Chemical communications 54, 948–951, doi:10.1039/c7cc08758f (2018).

65 Bartelt, S. M., Steinkühler, J., Dimova, R. & Wegner, S. V. Light-Guided Motility of a Minimal Synthetic Cell. Nano Letters 18, 7268–7274, doi:10.1021/acs.nanolett.8b03469 (2018).

66 Chung, I. et al. Spatial control of EGF receptor activation by reversible dimerization on living cells. Nature 464, 783–787, doi:10.1038/nature08827 (2010).

67 Van Haren, J., Adachi, L. S. & Wittmann, T. 211–234 (Springer US, 2020).

68 Paintdakhi, A. et al. Oufti: An integrated software package for high-accuracy, high-throughput quantitative microscopy analysis. Molecular microbiology 99, 767–777, doi:10.1111/mmi.13264 (2016).

69 Lew, M., Von Diezmann, A. R. S. & Moerner, W. E. Easy-DHPSF open-source software for three-dimensional localization of single molecules with precision beyond the optical diffraction limit. Protoc. Exch. (2013).

70 Yan, T., Richardson, C. J., Zhang, M. & Gahlmann, A. Computational correction of spatially variant optical aberrations in 3D single-molecule localization microscopy. Optics express 27, 12582–12599 (2019).

