## Supplementary Figures for "Dimerization of iLID Optogenetic Proteins Observed Using 3D Single-Molecule Tracking in Live Bacterial Cells"

**SUPPORTING INFORMATION**

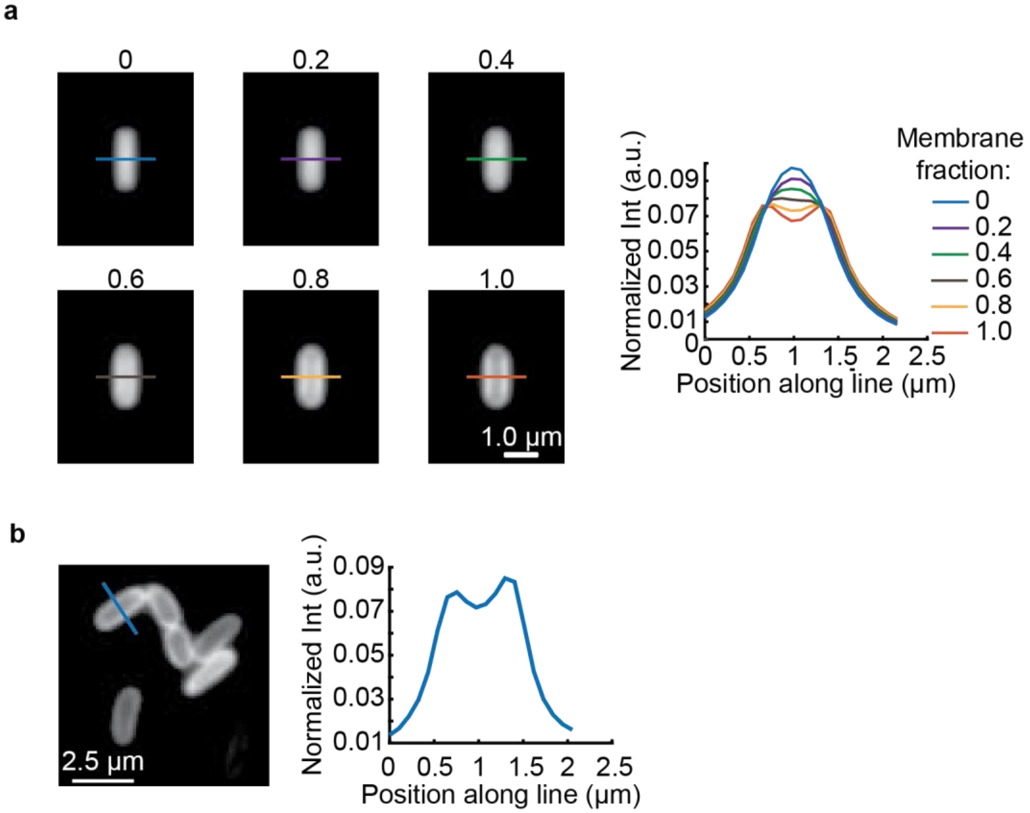


**SI Figure 1. (a)** Simulated diffraction-limited images of cells with increasing fractions of membrane-associated fluorophores show a clear shift in the fluorescence intensity line profiles across the midsection of cell. **(b)** Experimental fluorescence images and line profile observed for MA-mCherry-iLID qualitatively match that of simulated cells with 100% membrane-associated fluorophores.

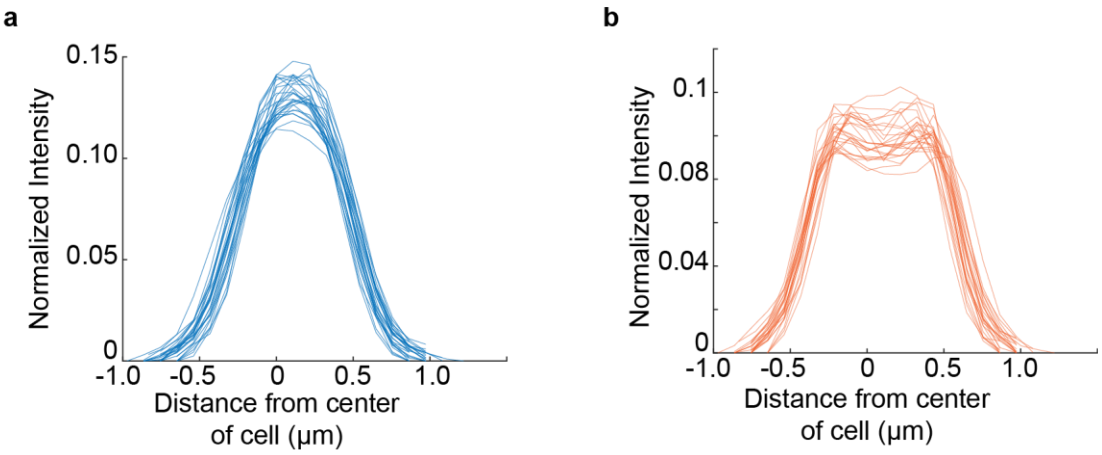


**SI Figure 2. Heterogeneity of spatial redistribution of $\mathbf{SspB_{nano}}$ upon 488 nm illumination.** (**a**) Overlay of normalized intensity line profiles prior to 488 nm illumination within a single field-of-view (n=27 cells). (**b**) Overlay of normalized intensity line profiles of the same cells as in (a) after 488 nm illumination.

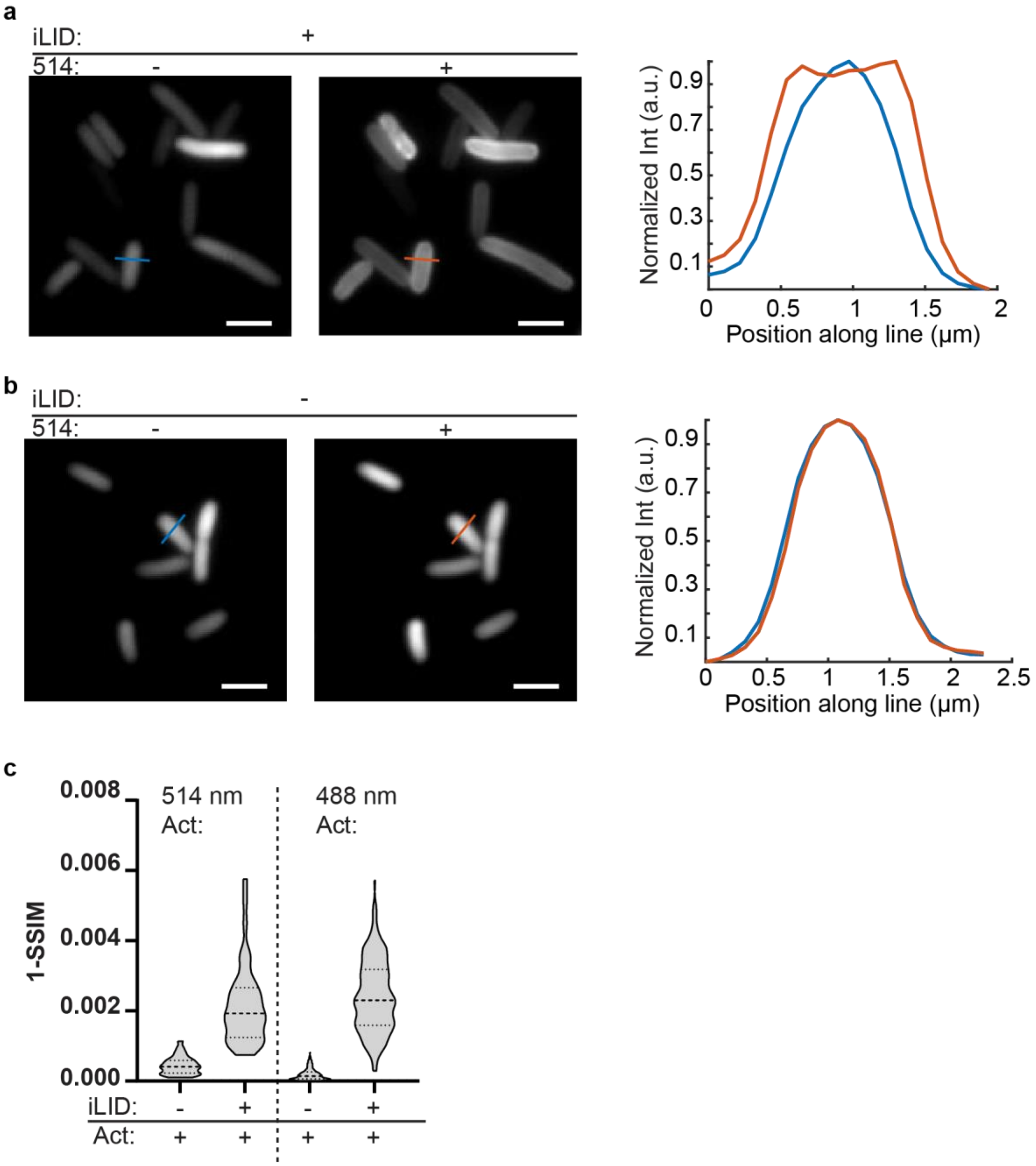


**SI Figure 3. Optical activation of the iLID: SspB$_{nano}$ interaction is possible with 514 nm light** **(a)** Spatial redistribution SspB$_{nano}$ is observed in cells expressing MA-iLID after 514 nm laser illumination. The fluorescence intensity line profile across the midsection of the cell changes from a Gaussian-like line shape to a characteristic double-peaked line shape. **(b)** Spatial redistribution of SspB$_{nano}$ is not observed after 514 nm light illumination in cells which do not express MA-iLID. The fluorescence intensity line profile across the midsection of the cell retains its Gaussian-like line shape **(c)** Image dissimilarity (1-SSIM) analysis performed on the full cell population before and after 514 nm light illumination (~1 W/cm$^2$). The image dissimilarity is comparable to cells illuminated with 488 nm light (~4 mW/ cm$^2$).

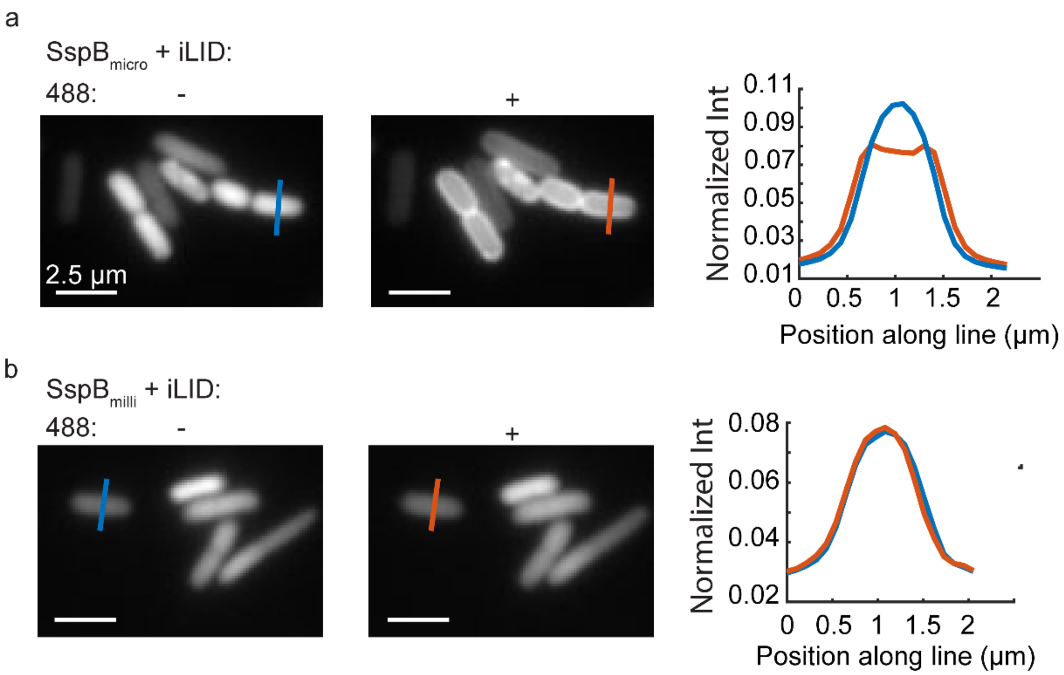


**SI Figure 4. Diffraction-limited images of SspB affinity mutants.** **(a)** Diffraction-limited images of SspB$_{micro}$ show robust redistribution of fluorescence from the cytosol to the membrane upon 488 nm light illumination. Normalized fluorescence line-profile information show distinct line shapes consistent with cytosolic and membrane-proximal fluorescence before and after 488 nm light illumination. **(b)** Diffraction-limited images of SspB$_{milli}$ do not indicate changes in fluorophore spatial distribution upon 488 nm light illumination.

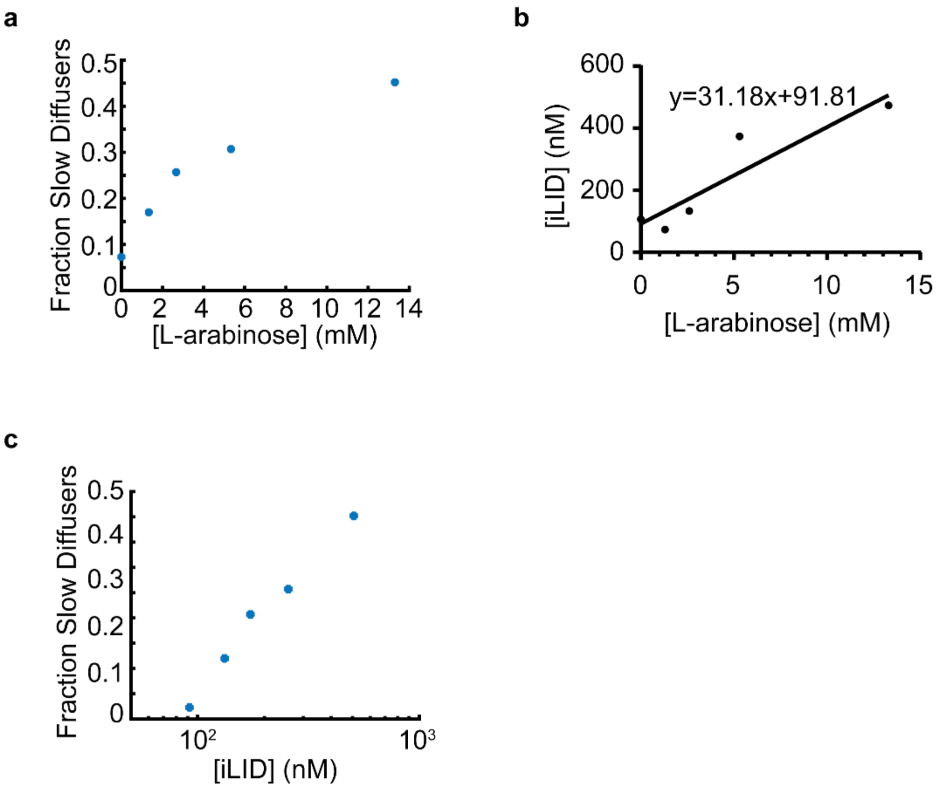


**SI Figure 5. The fraction of slow-diffusing, iLID-associated $SspB_{micro}$ is dependent on the expression level of MA-iLID. (a)** Fraction of slow diffusing $SspB_{micro}$ (estimated based on single-molecule tracking data) as a function of L-arabinose induction concentration. **(b)** Concentration of MA-mCherry-iLID as a function of L-arabinose concentration. The number of MA-iLID molecules was estimated by dividing initial total cell intensity of MA-mCherry-iLID expressing cells by the measured, average intensity of a single mCherry emitter. The data were fit to a line to estimate the MA-iLID concentrations at different L-arabinose induction concentrations. **(c)** Fraction of slowly diffusing $SspB_{micro}$ derived from single-molecule tracking data as a function of iLID concentration.

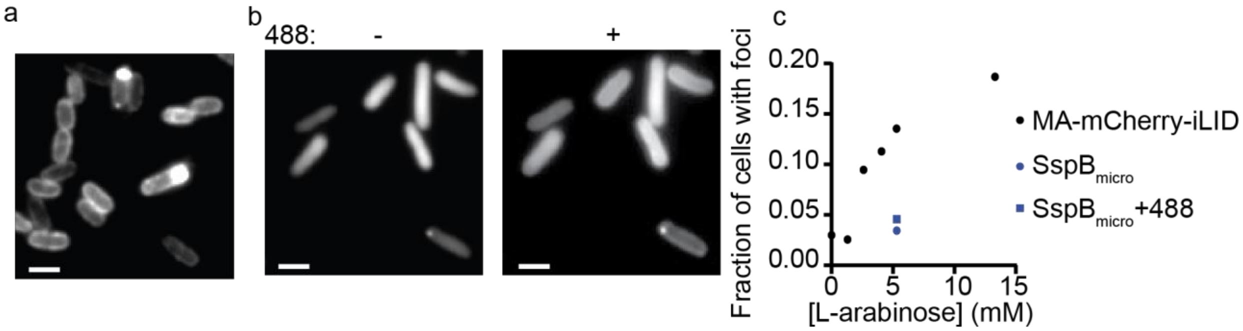


**SI Figure 6. Expression level-dependent fluorescent foci formation of MA-mCherry-iLID and $SspB_{micro}$. (a)** Diffraction-limited images of MA-mCherry-iLID expressing cells. The L-arabinose inducer concentration is 5.33 mM. Fluorescent foci formation is evident in a subset of cells. **(b)** Diffraction-limited images of $SspB_{micro}$ co-expressed with MA-iLID at the same L-arabinose concentration as in (a) before and after 488 nm illumination. **(c)** Quantification of the fraction cells with foci. 2 µm scale bar. (*N* = 618, 321, 81, 604, 445 cells for 0, 1.25, 2.5, 5.33, and 13.3 mM L-arabinose inducer concentrations, respectively; *N* = 88 cells for $SspB_{micro}$ + MA-iLID – pre and post 488 nm light illumination).

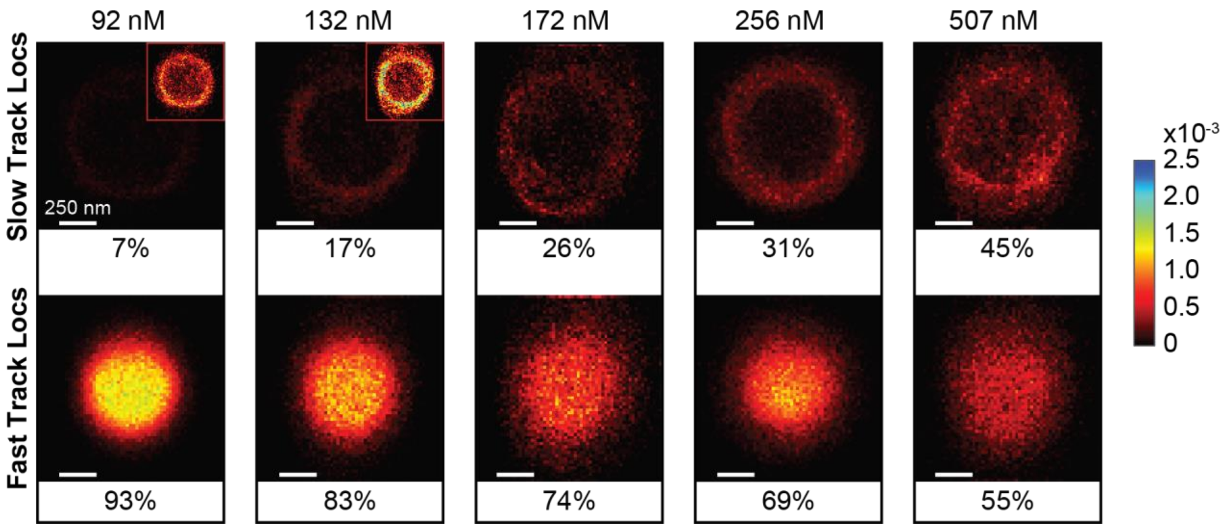


**SI Figure 7. Expression level of MA-iLID is directly correlated with fraction of slow diffusers and iLID-association.** 2D cross-sectional histograms of $SspB_{micro}$ single-molecule trajectories indicates increased probability of slow diffusion at the cell membrane as MA-iLID expression level is increased. Each histogram is normalized to the total number of molecules observed in that experiment. Inset: rescaled (unnormalized) histograms, shown for clarity. Inset histograms are rescaled versions of the full-size histogram, set to maximum values of $2.0\text{x}10^{-4}$ and to $3.0\text{x}10^{-4}$ for 92 nM and 132 nM expression levels, respectively.

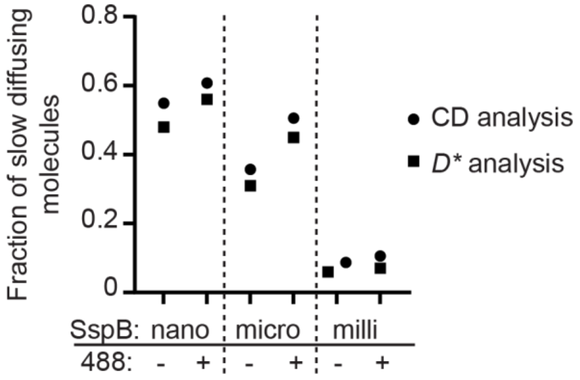


**SI Figure 8.** Fraction of slow diffusing molecules which stay bound for the duration of the trajectory in cumulative displacement (CD) analysis compared to fraction of slow diffusing molecules observed in apparent diffusion coefficient analysis (*D**).
